# Interplay between nucleoid-associated proteins and transcription factors in controlling specialized metabolism in *Streptomyces*

**DOI:** 10.1101/2021.04.15.440093

**Authors:** Xiafei Zhang, Sara N. Andres, Marie A. Elliot

## Abstract

Lsr2 is a small nucleoid-associated protein found throughout the actinobacteria. Lsr2 functions similarly to the well-studied H-NS, in that it preferentially binds AT-rich sequences and represses gene expression. In *Streptomyces venezuelae*, Lsr2 represses the expression of many specialized metabolic clusters, including the chloramphenicol antibiotic biosynthetic gene cluster, and deleting *lsr2* leads to significant upregulation of chloramphenicol cluster expression. We show here that Lsr2 likely exerts its repressive effects on the chloramphenicol cluster by polymerizing along the chromosome, and by bridging sites within and adjacent to the chloramphenicol cluster. CmlR is a known activator of the chloramphenicol cluster, but expression of its associated gene is not upregulated in an *lsr2* mutant strain. We demonstrate that CmlR is essential for chloramphenicol production, and further reveal that CmlR functions to ‘counter-silence’ Lsr2’s repressive effects by recruiting RNA polymerase and enhancing transcription, with RNA polymerase effectively clearing bound Lsr2 from the chloramphenicol cluster DNA. Our results provide insight into the interplay between opposing regulatory proteins that govern antibiotic production in *S. venezuelae*, which could be exploited to maximize the production of bioactive natural products in other systems.

**IMPORTANCE:** Specialized metabolic clusters in *Streptomyces* are the source of many clinically-prescribed antibiotics. However, many clusters are not expressed in the laboratory due to repression by the nucleoid-associated protein Lsr2. Understanding how Lsr2 represses cluster expression, and how repression can be alleviated, are key to accessing the metabolic potential of these bacteria. Using the chloramphenicol biosynthetic cluster from *Streptomyces venezuelae* as a model, we explored the mechanistic basis underlying Lsr2-mediated repression, and activation by the pathway-specific regulator CmlR. Lsr2 polymerized along the chromosome and bridged binding sites located within and outside of the cluster, promoting repression. Conversely, CmlR was essential for chloramphenicol production, and further functioned to counter-silence Lsr2 repression by recruiting RNA polymerase and promoting transcription, ultimately removing Lsr2 polymers from the chromosome. Manipulating the activity of both regulators led to >130× increase in chloramphenicol levels, suggesting that combinatorial regulatory strategies can be powerful tools for maximizing natural product yields.

## INTRODUCTION

*Streptomyces* species are renowned for their complex life cycle and their ability to produce a wide range of medically useful specialized metabolites, including over two-thirds of the antibiotics in clinical use today. Genome sequencing has revealed that most *Streptomyces* spp. encode 25-50 specialized metabolic clusters (1–3); however, the vast majority of their associated products have yet to be identified. Many of these clusters are poorly transcribed and consequently their resulting products have never been detected under laboratory conditions (4–6). These ‘cryptic’ and ‘silent’ clusters have the potential to produce an impressive array of novel antibiotics (1,7,8), and activating their expression is one of the keys to facilitating new antibiotic discovery.

In *Streptomyces*, specialized metabolic clusters are controlled by multiple factors. These include cluster-situated regulators (encoded within their cognate biosynthetic gene clusters) that govern metabolite synthesis by directly binding promoter regions in their associated cluster. Pleiotropic regulators have also been implicated in antibiotic control; these are usually encoded elsewhere on the chromosome and affect the expression of multiple biosynthetic clusters (9). In recent years, nucleoid-associated proteins have also been found to influence the expression of specialized metabolic clusters (4,10–12).

Historically, nucleoid-associated proteins function to promote chromosome organization; however, they can also impact activities like DNA replication, transcription, and chromosome segregation (13–15). H-NS (histone-like nucleoid-structuring protein) is one of the best-studied nucleoid-associated proteins. It is, however, only found in a subset of Gram-negative bacteria, where it preferentially binds and spreads along and/or bridges distal high AT-content DNA, compacting the chromosome and/or silencing gene expression (14,16–19). The resulting DNA filaments and/or DNA bridges formed by H-NS have the potential to affect gene expression by trapping RNA polymerase and repressing transcription, or by excluding RNA polymerase from promoter regions.

In the streptomycetes, H-NS-like proteins play important roles in regulating antibiotic production. The H-NS-equivalent protein in these bacteria is termed Lsr2, and it is conserved throughout the actinobacteria (15,20). Like H-NS, Lsr2 is a global repressor that preferentially binds high AT-content DNA (4,20,21), and based on work with the mycobacterial protein, is predicted to silence gene transcription by bridging or oligomerizing along the DNA (16,17,20). Deleting *lsr2* in *Streptomyces venezuelae* leads to significantly upregulated gene expression in a majority of specialized metabolic biosynthetic clusters, including many otherwise ‘cryptic’ clusters that are not expressed in a wild type background (4). This suggests that Lsr2 functions to broadly repress specialized metabolism in *Streptomyces* species.

To better understand how Lsr2 repression is both exerted and alleviated in the streptomycetes, we focussed our attention on the chloramphenicol biosynthetic cluster. Previous work revealed that loss of Lsr2 results in a dramatic increase in the expression of the chloramphenicol biosynthetic genes, and this effect seems to be a direct one, as an Lsr2 binding site was identified within the gene cluster (4) (**Fig. 1**). The chloramphenicol biosynthetic cluster comprises 16 genes (*sven0913-sven0928*), with *sven0913/cmlR* encoding a pathway-specific transcriptional activator (22). Here, we show that Lsr2 binding to the cluster-internal site, and to an upstream adjacent sequence, leads to Lsr2 polymerization along the DNA, and can promote bridging between these two regions. This binding activity limits chloramphenicol production, presumably through the repression of cluster transcription. Lsr2 repression can be relieved through the action of CmlR, which functions as a counter-silencer of Lsr2 activity and is essential for chloramphenicol production. CmlR appears to exert its activity not by competing with Lsr2 for binding, but instead by promoting cluster transcription, where the action of RNA polymerase serves to clear Lsr2 from the DNA, alleviating cluster repression.

**Fig 1:**
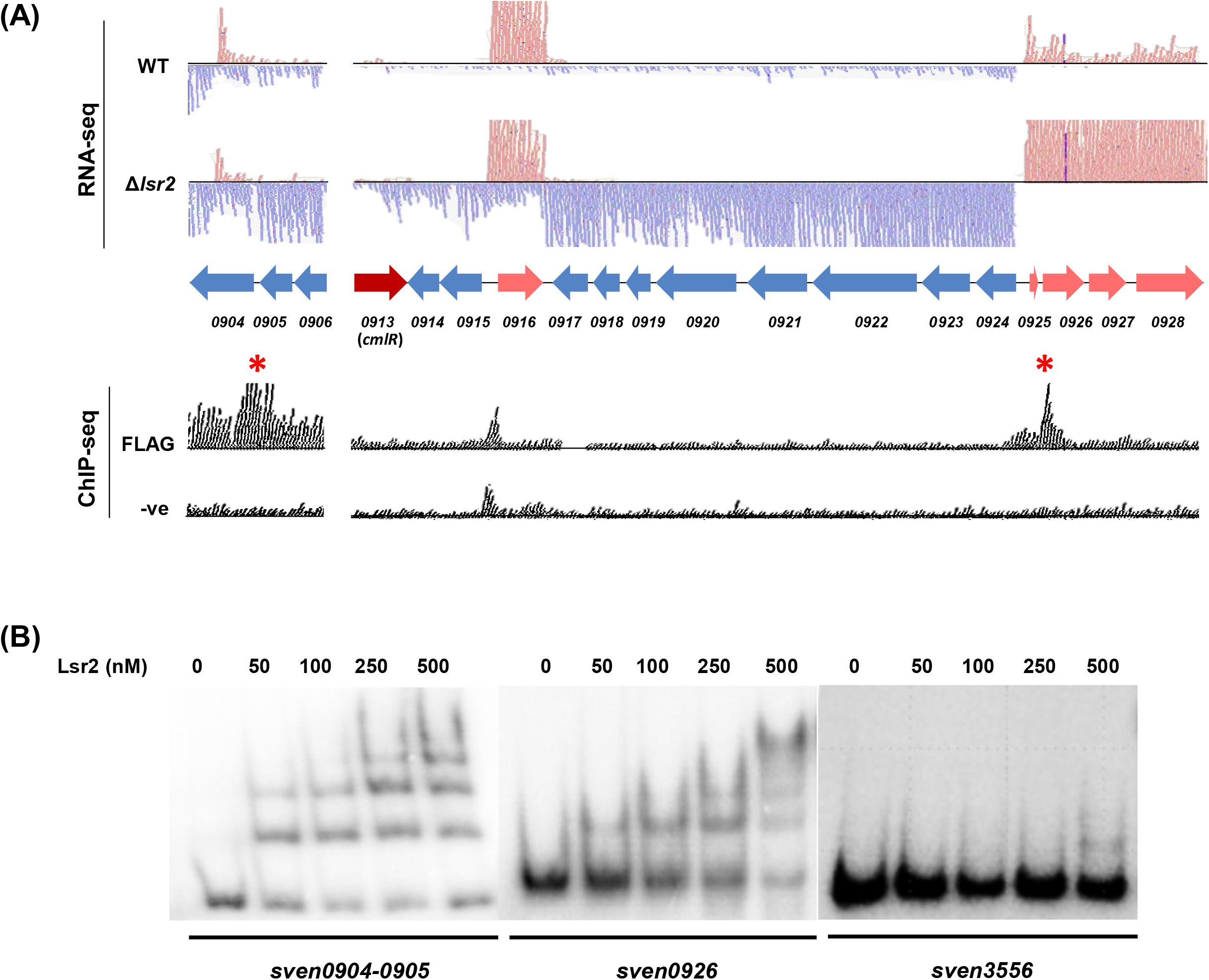
Lsr2 binding sites and effect on transcription of the chloramphenicol biosynthetic cluster. **(A)** Top panels: RNA-seq analysis of gene expression within and upstream of the chloramphenicol biosynthetic cluster in wild type and *lsr2* mutant strains. Blue reads (and gene arrows) map to the reverse strand, and pink reads (and gene arrows) map to the forward strand. Bottom panels: ChIP-seq analysis of Lsr2 binding sites (using a FLAG-tagged Lsr2 variant), alongside a negative control (expressing untagged Lsr2). Red asterisks indicate statistically significant Lsr2 binding sites. **(B)** EMSAs probing Lsr2 binding to sites within and adjacent to the chloramphenicol biosynthetic cluster. Increasing concentrations of Lsr2 (0-500 nM) were combined with 1 nM of labelled *sven0904-05* (upstream/adjacent), *sven0926* (internal), or *sven3556* (negative control) probes. The results presented are representative of two independent biological replicates.

## RESULTS

### Antibiotic production is impacted by Lsr2 binding to sites adjacent to the chloramphenicol cluster

Lsr2 represses the expression of the majority of genes in the chloramphenicol biosynthetic cluster (4) (**Fig. 1A**). Intriguingly, the only Lsr2 binding site within the cluster was in the coding sequence of a gene (*sven0926*) located at the 3′ end of the cluster (4) (**Fig. 1A**). We revisited our chromatin-immunoprecipitation sequencing (ChIP-seq) data (4) and noted that there was a second Lsr2 binding site upstream of the cluster, spanning genes *sven0904-0905*, where *sven0904* is predicted to encode a solute binding transport lipoprotein, and *sven0905* is predicted to encode a short chain oxidoreductase (**Fig. 1A**). We first set out to validate Lsr2 binding to both internal and upstream sites using EMSAs. We found Lsr2 had a much higher affinity for *sven0904-0905* and *sven0926* probes than for a negative control sequence (within *sven3556*, which was not bound by Lsr2 in our previous ChIP-seq experiments), confirming the specific binding of Lsr2 to these sites within and adjacent to the chloramphenicol cluster (**Fig. 1B**).

Given the functional similarity shared by Lsr2 and H-NS, we hypothesized that Lsr2 may exert its repressive effects in a manner analogous to H-NS, either by polymerizing along the DNA and/or bridging distant DNA regions. We considered three mechanisms by which Lsr2 could repress transcription of the chloramphenicol cluster: (1) Lsr2 could bind within *sven0926* and polymerize along the chromosome, repressing expression of the flanking gene clusters; (2) Lsr2 could bind to both *sven0904-0905* and *sven0926* sites, and interact to bridge these sequences and alter the structure of the intervening DNA, or (3) Lsr2 could both polymerize along the DNA and bridge these disparate sequences. We expected that if Lsr2 repressed transcription of the chloramphenicol cluster by only polymerizing from the *sven0926* binding site, then the *sven0904-0905* binding site would be dispensable for Lsr2 repression, and this region would have no effect on chloramphenicol production. If, however, Lsr2 repression was mediated by bridging these two sites (*sven0926* and *sven0904-0905*), or both polymerizing along the DNA and bridging these two regions, then deleting the upstream/cluster-adjacent binding site would relieve cluster repression, and yield increased chloramphenicol levels relative to the wild type strain.

To probe these different scenarios, we compared chloramphenicol production by wild type and Δ*lsr2* strains, alongside a Δ*0904-0905* and double Δ*lsr2*Δ*0904-0905* mutant strain using liquid chromatography–mass spectrometry (LC-MS). LC-MS analyses revealed that, relative to wild type, deleting *sven0904-0905* led to a significant increase (~4-fold) in chloramphenicol production, while deleting both *lsr2* and *sven0904-0905* led to a ~13-fold increase in chloramphenicol production, which was similar to the production levels of the Δ*lsr2* mutant alone (~11-fold) (**Fig. 2**).

**Fig. 2:**
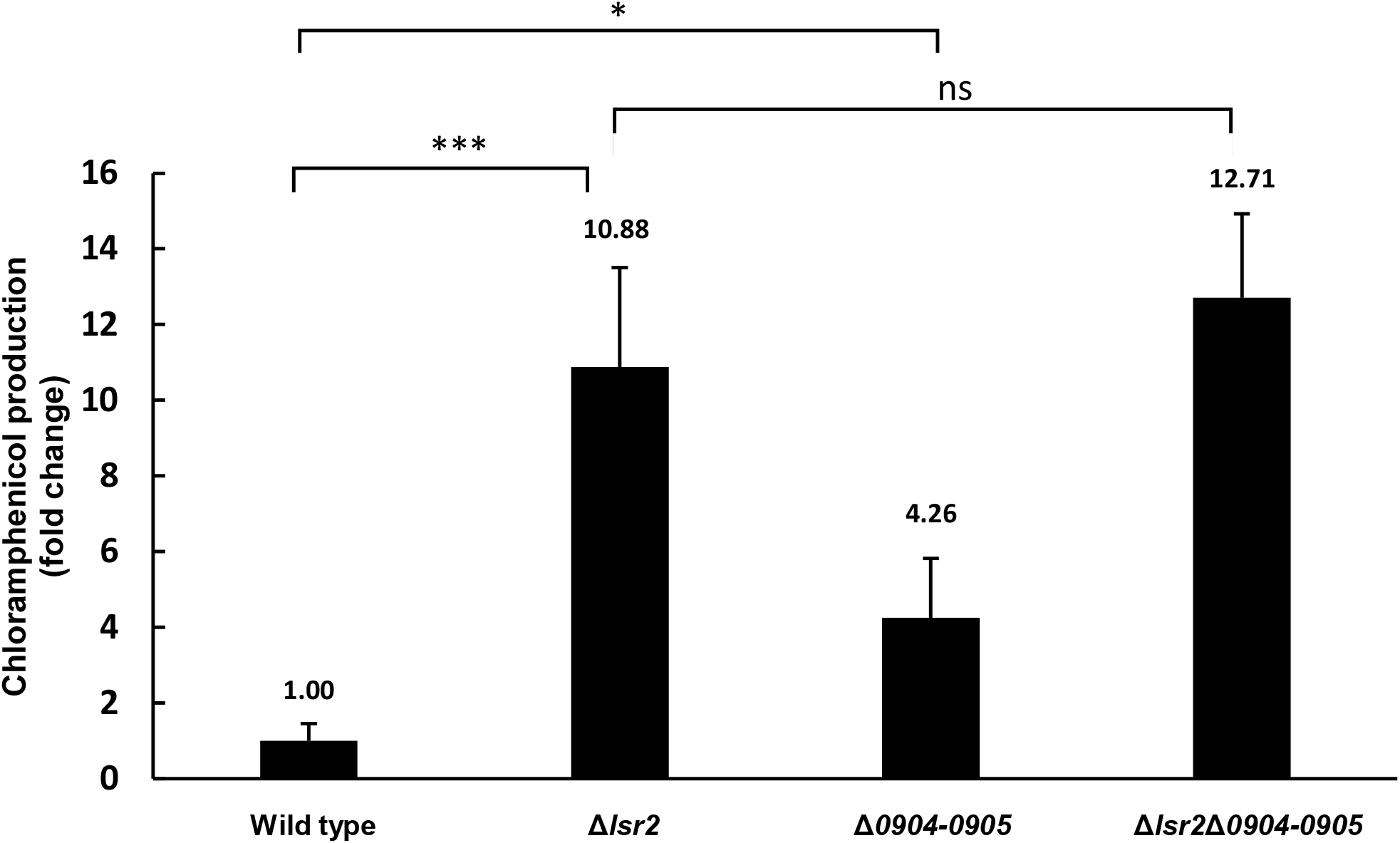
Deleting *sven0904-0905* from the chromosome increased chloramphenicol production. *sven0904-0905* were deleted in wild type and Δ*lsr2* backgrounds, and LC-MS analyses were performed on the resulting strains after 2 days growth in liquid culture, to quantify changes in chloramphenicol production relative to wild type. Error bars represent standard deviation for three independent biological replicates. * indicates p<0.05; *** indicates p<0.005; ns indicates no significant difference.

These results were consistent with a possible role for the *sven0904-0905* site in repressing chloramphenicol production through Lsr2 bridging between this site and the internal binding site. It was, however, formally possible that the products of these two upstream genes negatively influenced chloramphenicol production. To test this second possibility, we sought to complement the *sven0904-0905* mutant strains by cloning the operon containing wild type *sven0904-0905* into the integrating plasmid vector pMS82. We reasoned that reintroducing these genes on a plasmid vector that integrated at an independent site in the chromosome should restore wild type levels of chloramphenicol production if their products were important for antibiotic production, whereas no complementation of the mutant phenotype was expected if the locus position was critical for cluster repression. We introduced the complementation construct into the mutant strains, alongside the empty plasmid as a control (in both mutants and wild type), and assessed chloramphenicol production by these different strains. Complementing the mutants (Δ*0904-0905* and Δ*lsr2*Δ*0904-0905*) with the *sven0904-0906* operon failed to restore production levels to that of the empty plasmid-containing wild type and Δ*lsr2* strains (**Supplementary Fig. 1**). This suggested that the position of the *sven0904-0905* locus on the chromosome (and its associated Lsr2-binding site) – and not the function of the SVEN0904 and 0905 gene products – may be important for controlling chloramphenicol production.

### Lsr2 binding leads to polymerization along the DNA and bridging between sites upstream and within the chloramphenicol cluster

To explore the potential bridging capabilities of Lsr2, we employed atomic force microscopy (AFM). The two chloramphenicol cluster-associated Lsr2 binding sites are separated by 24 kb, which would be larger than ideal for use in AFM experiments. We initially opted to bring these two binding sites closer together, such that there was ~1 kb separating each core binding site (giving a total DNA fragment length of 2919 bp). Lsr2 was then added, and the resulting products were visualized. If DNA bridging was the sole mechanism by which Lsr2 exerted its regulatory activity, we expected to see a loop formed between the Lsr2 binding sites at either end of the DNA fragment. However, we failed to detect any loop structures, and instead observed only DNA molecules that had been coated and compacted by Lsr2, suggesting that Lsr2 could polymerize along the DNA under these *in vitro* conditions.

To better assess the bridging potential of Lsr2, we added an extra 1 kb of sequence between the two Lsr2 binding sites, to give a DNA fragment of ~4 kb in size. Using AFM, we compared the length of the DNA alone, with that of DNA mixed with Lsr2. For the DNA-alone experiments, we needed to supplement the binding buffer with Ni^2+^ to facilitate DNA adherence to the mica slide used for the AFM experiments; Ni^2+^ was not added to the Lsr2-containing samples, as it disrupted DNA binding by Lsr2. For the DNA-alone controls, we observed linear DNA molecules (**Fig. 3A,B**), with an average length of 1273.7 nm (*n*=71) (**Fig. 3B,C**), consistent with the expected length of 1200 nm for a 4 kb DNA molecule. In the presence of 250 nM of Lsr2, looped molecules were identified alongside linear-appearing DNA-Lsr2 complexes (*n*=54) (**Fig. 3A,B**). For the linear-appearing DNA-Lsr2 complexes, Lsr2 polymerization was apparent at one end of the DNA, but no obvious bridging was observed. In contrast, loop structures appeared to result from Lsr2 bridging the two distal regions. Notably, Lsr2 polymerization was also typically observed at each bridging site, where the loop appeared to have been ‘zipped up’ (**Fig. 3A**). The lengths of both the looped and linear DNA-Lsr2 complexes were measured in the presence of 250 nM of Lsr2, and the mean value was found to be 845.7 nm (*n*=54) (**Fig. 3B,C**). To ensure these changes in DNA structure and length stemmed from specific Lsr2 binding and oligomerization, and not simply DNA folding back on itself, the height of the observed DNA alone molecules and Lsr2-bound regions were measured; it was expected that Lsr2 binding to DNA would result in a minimum 3-fold increase in height. The mean values of the height of DNA alone and Lsr2-bound regions were 0.23 nm (*n=*71) and 1.15 nm (*n=*36), respectively (**Fig. 3B,C**). To further confirm that these DNA structures were the result of specific Lsr2 binding, equivalent experiments were performed using a 4.5 kb DNA fragment that lacked Lsr2 binding sites, based on our previous ChIP-seq analyses (4). As expected, DNA alone adopted a linear configuration. However, under the conditions used for Lsr2 binding, we consistently failed to detect any DNA, suggesting that Lsr2 was unable to specifically associate with this DNA fragment and tether the DNA to the slide (**Supplementary Fig. 2**). In all, the AFM results suggested that Lsr2 could polymerize along the DNA, and had the capacity to bridge disparately positioned sites (at least 4 kb apart) and polymerize towards each binding site. These collective actions may serve to down-regulate chloramphenicol production by limiting RNA polymerase access/activity within the chloramphenicol biosynthetic cluster in *S. venezuelae*.

**Fig. 3:**
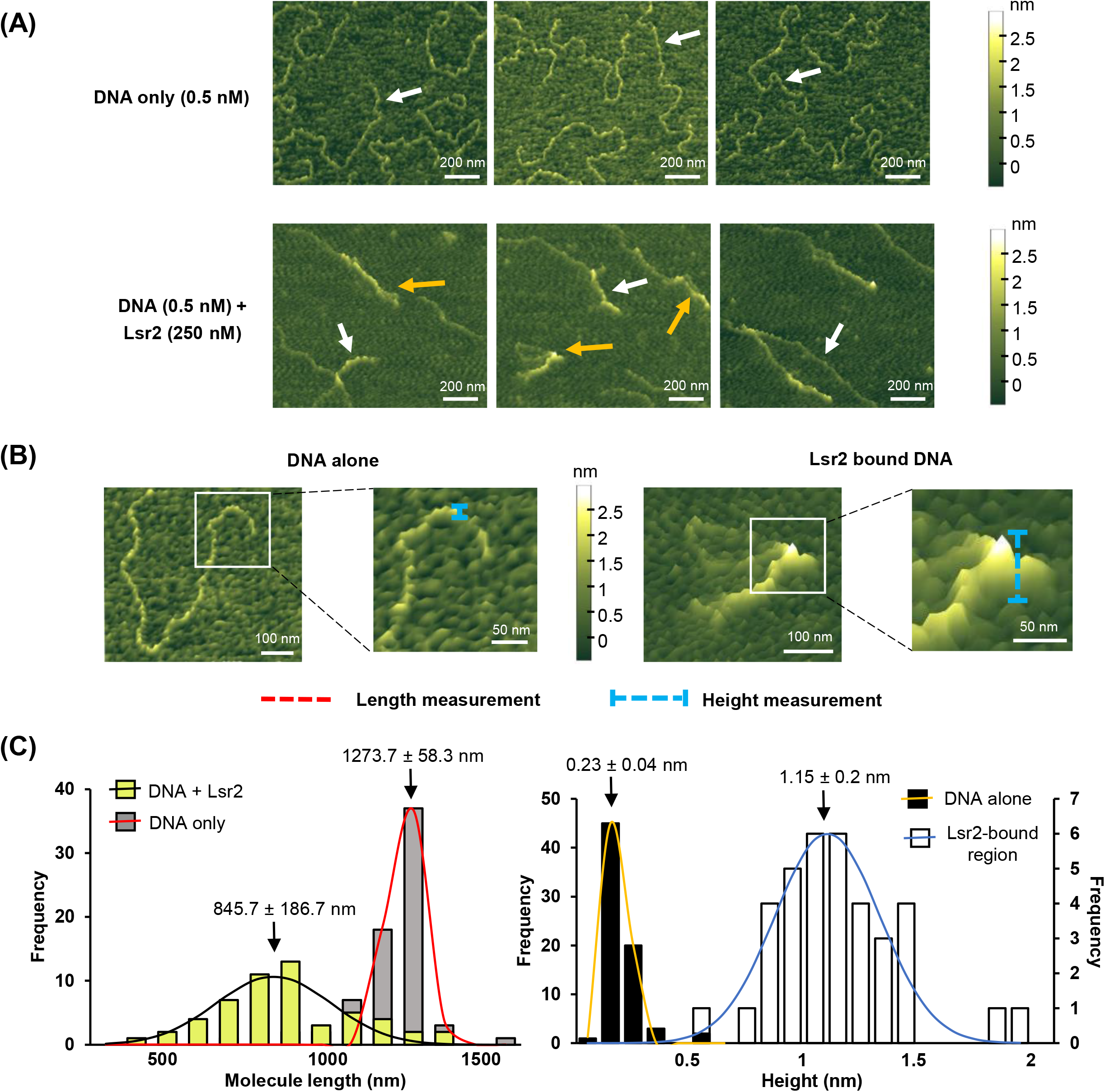
Lsr2 binds specific target sequences and can both form polymers along the DNA, and bridge binding sites. **(A)** Top: atomic force microscopy (AFM) images of engineered (target) DNA molecules with two Lsr2 binding sites on either end. Bottom: AFM images of 0.5 nM target DNA + 250 nM Lsr2. White arrows indicate linear DNA molecules; orange arrows indicate looped structures. **(B)** Illustration of how length and height measurements of DNA alone and Lsr2-bound regions were taken. **(C)** Left: Frequency distribution of length of Lsr2-bound/unbound DNA molecules. *n=*71 for DNA only and 54 for DNA + Lsr2. Right: Frequency distribution of mean height of DNA alone (frequency axis on the left) and Lsr2-bound regions (frequency axis on the right). *n=*71 for DNA alone and 36 for Lsr2-bound regions. Data are presented as mean ± standard deviation, calculated from nonlinear Gaussian fit.

### The pathway-specific regulator CmlR is essential for chloramphenicol production

We next set out to understand how the cluster-situated regulator CmlR impacted chloramphenicol production. Our previous RNA sequencing results had revealed that the expression of most genes in the chloramphenicol biosynthetic cluster was significantly increased in the absence of Lsr2. A notable exception, however, was *cmlR* (*sven0913*), whose transcript levels were consistent in both wild type and *lsr2* mutant strains (**Fig. 1A**). Consequently, we wondered whether CmlR might function simply to relieve Lsr2 repression, and if it was dispensable for cluster expression in the absence of Lsr2.

To test this hypothesis, we sought to determine the relative importance of CmlR in wild type and *lsr2* mutant strains of *S. venezuelae*. We created strains in which *cmlR* was deleted from the chromosome, and where it was overexpressed from a strong, constitutive (*ermE\**) promoter on an integrating plasmid in both wild type and *lsr2* mutant strains. We then tested chloramphenicol production levels in these different strains using LC-MS analyses. In these experiments, we found that deleting *lsr2* led to an ~8-fold increase in chloramphenicol production relative to wild type, and that deleting *cmlR* abolished chloramphenicol production in all strains. This suggested CmlR was critical for chloramphenicol biosynthesis beyond simply relieving Lsr2 repression. Consistent with this observation, we found that overexpressing *cmlR* led to a massive increase in chloramphenicol production in wild type strains (102-fold increase relative to plasmid-alone controls), while overexpressing CmlR in the absence of Lsr2 led to even higher chloramphenicol levels (134-fold increase) (**Fig. 4**). These results suggested that CmlR activity was essential for stimulating chloramphenicol production.

**Fig. 4:**
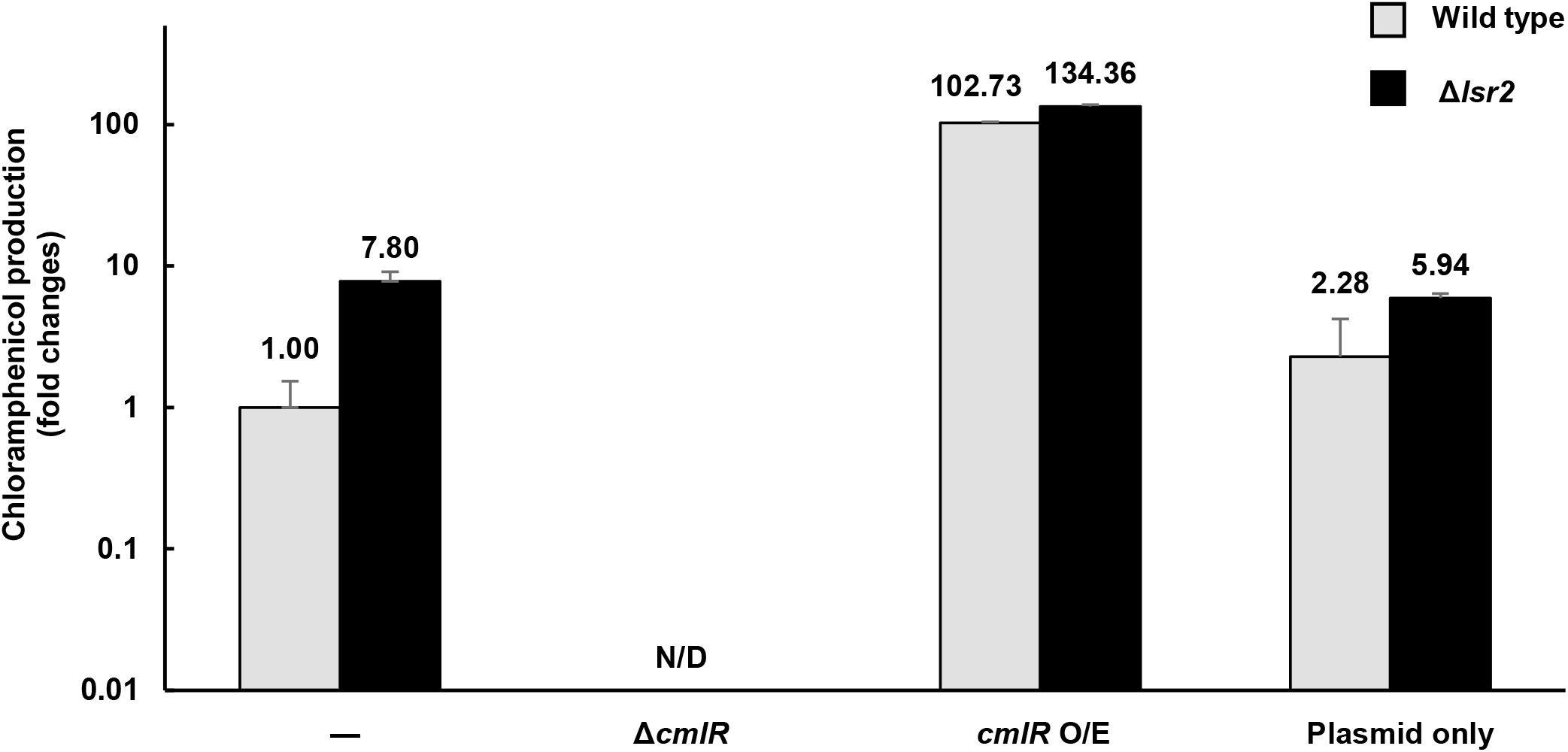
CmlR is required for chloramphenicol production in *S. venezuelae*. LC-MS analyses of changes in chloramphenicol production, relative to wild type, are plotted on a logarithmic graph. Grey bars: wild type background. Black bars: Δ*lsr2* background. N/D: not detected. Error bars represent standard deviation for two independent biological replicates.

### CmlR binds to a divergent promoter region in the chloramphenicol biosynthetic cluster

To begin to understand how CmlR exerted its regulatory effects within the chloramphenicol cluster, we examined its DNA binding capabilities. CmlR shares 44% amino acid sequence identity with StrR (22), which is the pathway-specific activator of the streptomycin biosynthetic gene cluster in *Streptomyces griseus* (23). The StrR target sequence is well-established (5’-GTTCGActG(N)_11_CagTcGAAc-3’) (23), and so we searched for similar sequences in the intergenic/promoter-containing regions of the chloramphenicol cluster. We identified a potential binding site for CmlR between the *sven0924* and *sven0925* promoters, upstream of the Lsr2 binding site within *sven0926* (**Fig. 5A**). To test whether CmlR specifically bound this sequence, we conducted EMSAs using the predicted binding site as a probe. We found that CmlR directly bound the promoter region with high affinity (**Fig. 5B**). We confirmed binding specificity using the promoter of a gene outside of the chloramphenicol cluster (*sven5133*); there was no binding to this DNA fragment when using equivalent concentrations of CmlR (**Fig. 5B**). This implied that CmlR specifically bound a site between the promoters driving the *sven0924-* and *sven0925-*associated operons.

**Fig. 5:**
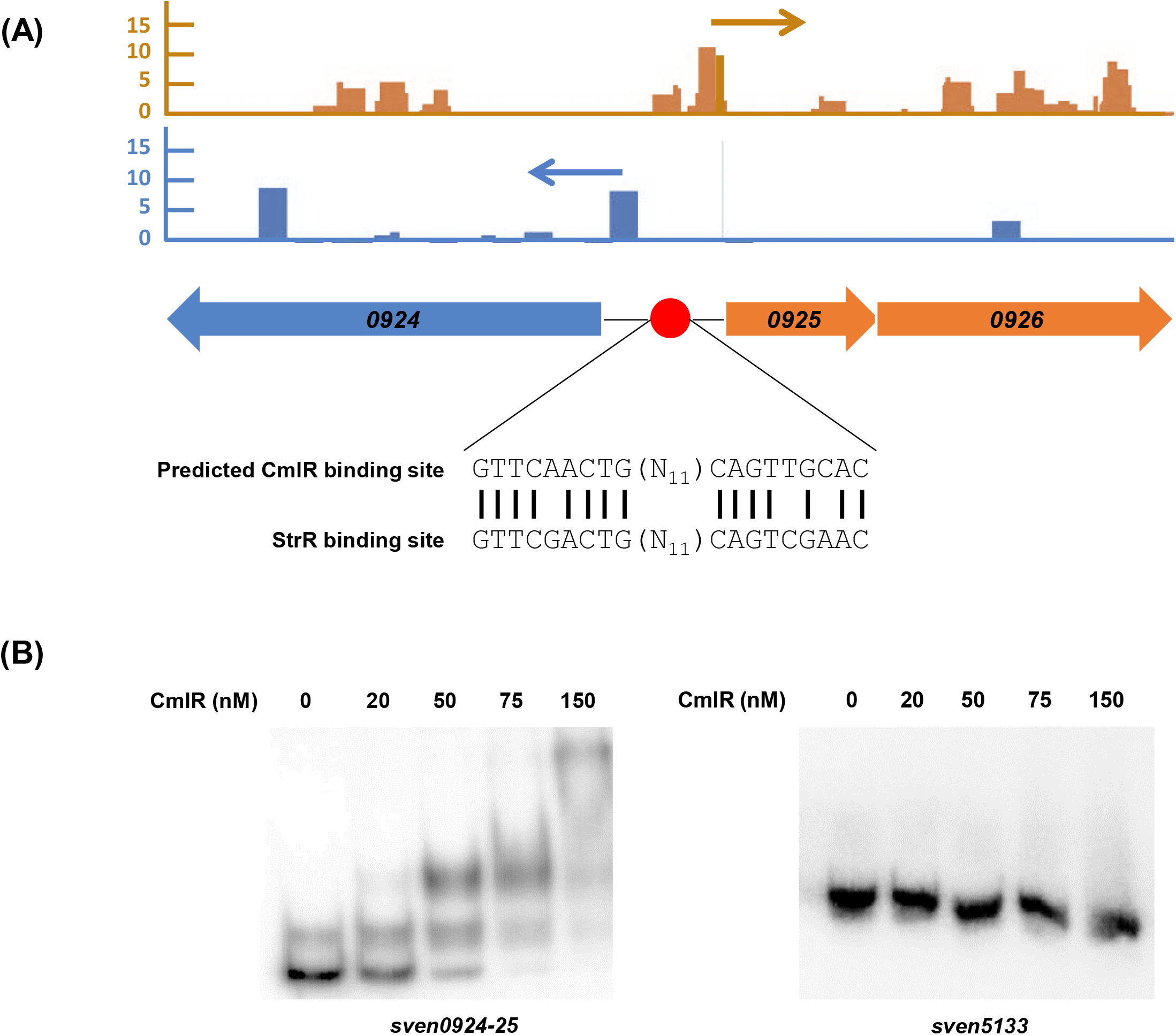
CmlR binds promoter regions within the chloramphenicol biosynthetic cluster. **(A)** Top: transcription start sites mapped for the *sven0924* and *sven0925* operons, as determined using differential RNA sequencing. Middle: schematic diagram showing the predicted CmlR binding site at the divergent promoter region between *sven0924* and *sven0925.* Red circle: CmlR. Blue reads (and blue arrows) map to the reverse strand; orange reads (and orange arrows) map to the forward strand. **(B)** EMSA using 1 nM of labelled *sven0924-0925* or *sven5133* (negative control) promoter regions as probe, together with increasing concentrations (0-150 nM) of purified CmlR. Results are representative of two independent mobility shift assays.

### CmlR alleviates Lsr2 repression within the chloramphenicol biosynthetic cluster

Given the relative proximity of the CmlR and Lsr2 binding sites within the chloramphenicol cluster, and that CmlR overexpression appeared to overcome Lsr2-mediated repression of cluster expression, we wanted to determine whether CmlR could act to ‘counter-silence’ the repressive effects of Lsr2. To address this possibility, we tested whether overexpressing CmlR reduced Lsr2 binding within the chloramphenicol cluster. We introduced our Lsr2-FLAG-tagged expression construct into the double *lsr2, cmlR* mutant strain, and into an *lsr2* mutant strain overexpressing CmlR. Using ChIP to capture DNA sequences bound by Lsr2-FLAG, we then used quantitative PCR (qPCR) to compare the relative amount of target DNA (*sven0926*) bound by Lsr2 in strains lacking or overexpressing CmlR. To ensure that any CmlR-mediated effects were specific to Lsr2 binding within the chloramphenicol cluster, we also assessed Lsr2 binding to another validated Lsr2-binding site positioned outside of the chloramphenicol cluster (*sven6264*), alongside negative controls sequences not bound by Lsr2 (based on previous ChIP experiments) (4).

Overexpressing CmlR reduced the levels of *sven0926* bound by Lsr2 by 40%, while deleting *cmlR* resulted in >100% increase in *sven0926* bound by Lsr2, relative to that bound by Lsr2 in the presence of wild type levels of CmlR (**Fig. 6**). Overexpressing and deleting *cmlR* had no obvious effects on either the abundance of the external Lsr2 target *sven6264*, or the negative control sequence. Taken together, these findings indicated that CmlR activity could influence Lsr2 binding within the chloramphenicol biosynthetic cluster, and in doing so, had the potential to counter the repressive effects of Lsr2.

**Fig. 6:**
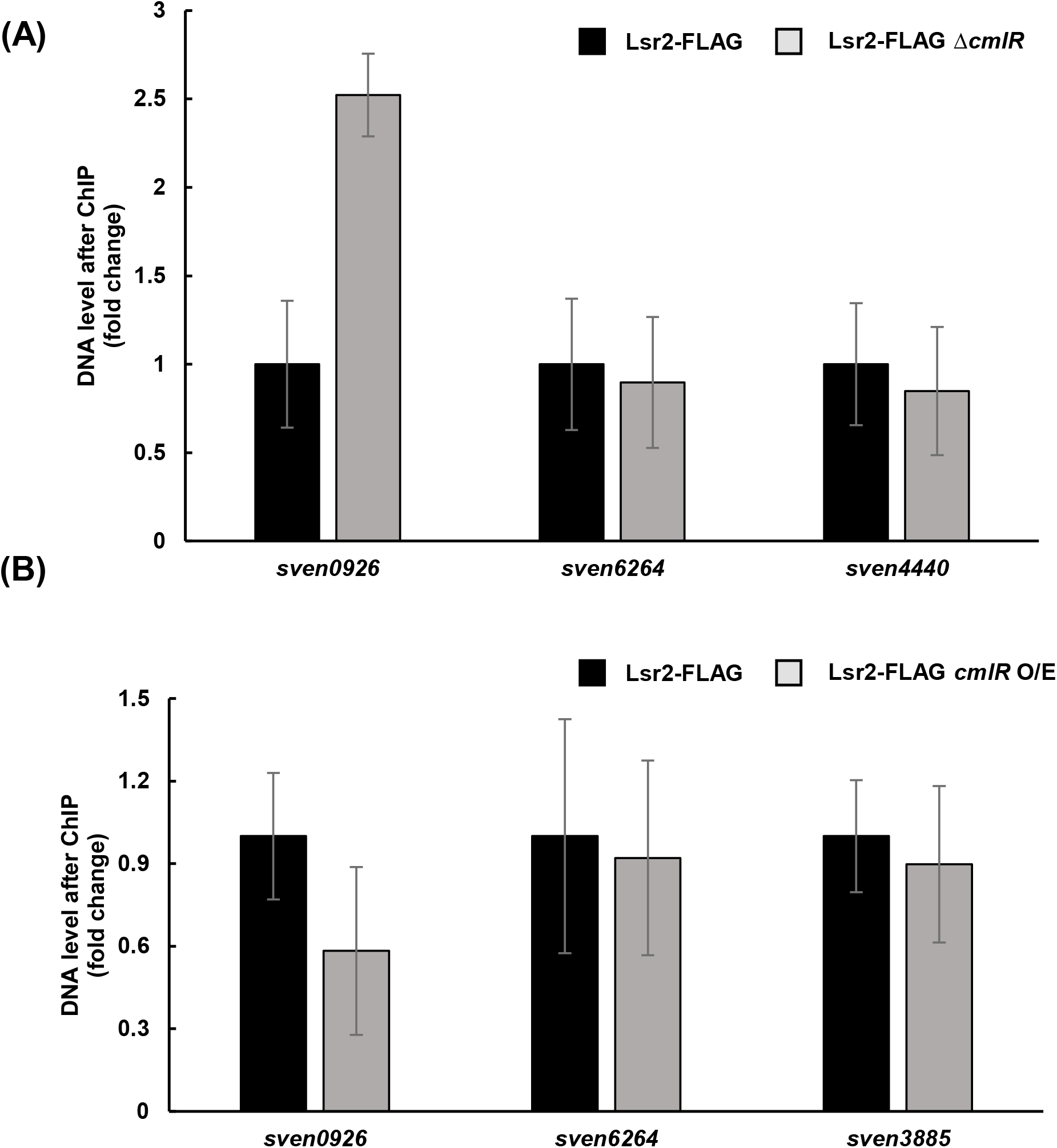
CmlR levels affect Lsr2 binding. **(A)** ChIP-qPCR quantification of the relative abundance of *sven0926, sven6264* (Lsr2-binding site positioned outside of the chloramphenicol cluster) and *sven4440* (negative control; not bound by Lsr2 in ChIP experiments) bound by Lsr2, in a strain with and without *cmlR* (black and grey bars, respectively). **(B)** qPCR analysis of ChIP DNA samples, quantifying the relative abundance of *sven0926*, *sven6264* and *sven3885* (negative control; not bound by Lsr2 in ChIP experiments) in a strain with wild type *cmlR* (black bars), versus a *cmlR* overexpression (O/E) strain (grey bars). For both (A) and (B), error bars represent standard error of the mean, for technical triplicate, and biological duplicate samples.

### CmlR alleviates Lsr2 repression by enhancing transcription

How CmlR impacted Lsr2 binding was not immediately obvious. We hypothesized that CmlR functioned to recruit RNA polymerase, and that the act of transcription disrupted the Lsr2 polymers/bridges, thus relieving Lsr2 repression. To test this possibility, we assessed whether inhibiting transcription affected Lsr2 binding, taking advantage of the fact that RNA polymerase (and correspondingly transcript elongation) could be inhibited by the antibiotic rifampicin (24,25).

Using a strain expressing the FLAG-tagged Lsr2 protein and overexpressing CmlR, we performed ChIP experiments after a 10 min exposure to rifampicin. In parallel, ChIP experiments were done using an untreated control strain. We quantified and compared the levels of *sven0926* bound by Lsr2, both with and without rifampicin treatment, using qPCR. We knew that overexpressing CmlR reduced the levels of *sven0926* bound by Lsr2 (**Fig. 7**). Thus we hypothesized that if CmlR alleviated Lsr2 binding and cluster repression by recruiting RNA polymerase and enhancing transcription, then inhibiting RNA polymerase activity would lead to increased Lsr2 binding to *sven0926*. We found that adding rifampicin to a CmlR overexpressing strain, led to a >500% increase in the amount of *sven0926* bound by Lsr2, relative to untreated controls. This suggested that CmlR relieved Lsr2 silencing by recruiting RNA polymerase, and the resulting increase in transcription served to remove Lsr2 polymers from the chromosome, and/or disrupt Lsr2 bridges.

**Fig. 7:**
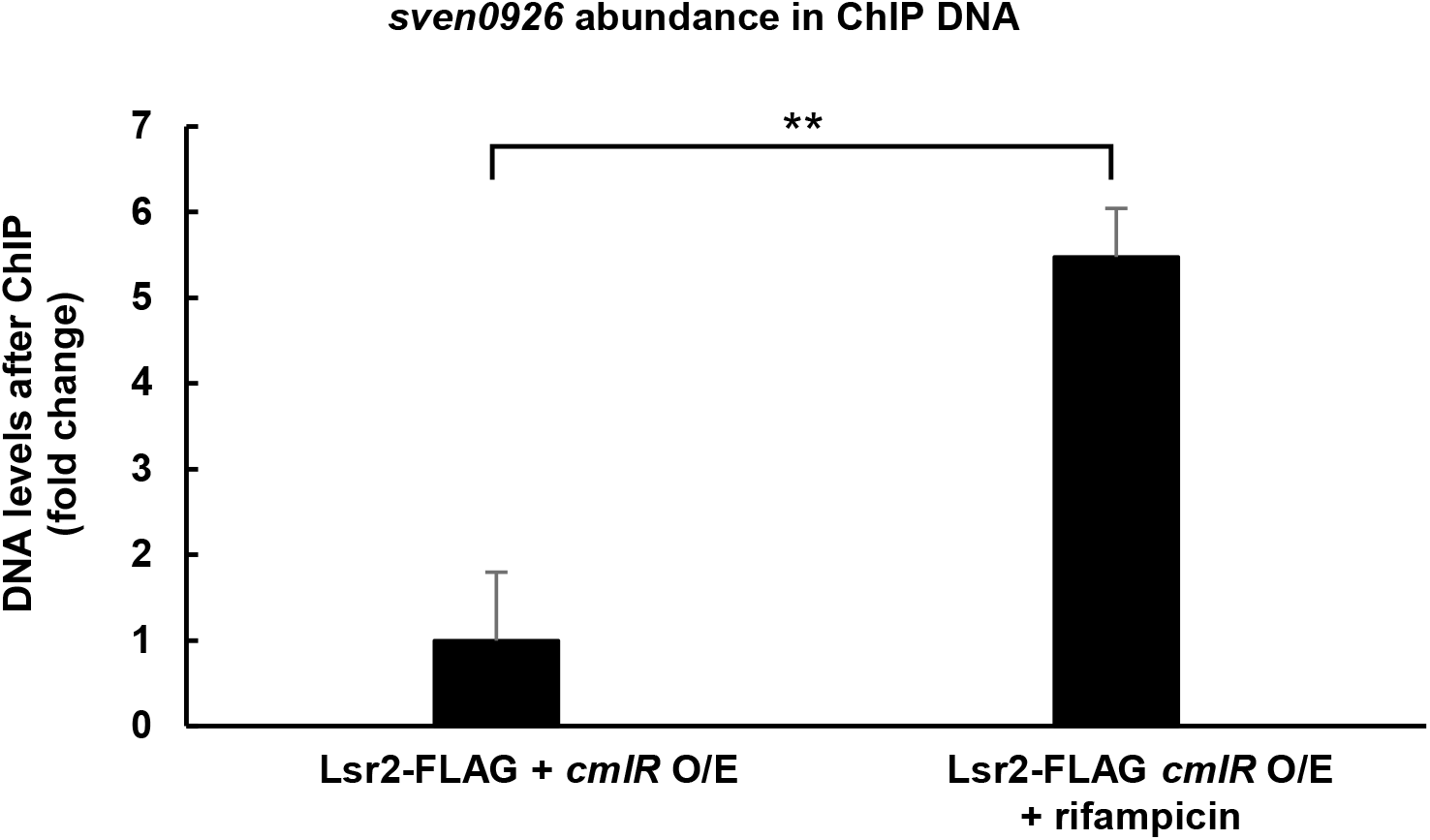
Inhibiting transcription enhances Lsr2 binding to its *sven0926* target site. The relative abundance (fold change) of Lsr2-targeted *sven0926* in rifampicin-treated (and untreated) *cmlR* overexpression strains was compared using qPCR, with ChIP-DNA samples as template. Error bars represent the standard error of the mean, for technical triplicates and biological duplicates. ** indicates p<0.01.

To further test the proposed mechanism of CmlR-mediated counter-silencing of Lsr2 activity, we explored the effects of CmlR using a simplified system in which Lsr2 repression could only be exerted by polymerizing along the chromosome. We employed a transcriptional reporter system, and fused two distinct promoter constructs to the *gusA* (β-glucuronidase-encoding) reporter gene (**Supplementary Fig. 3A**). The first contained the CmlR binding site and promoter for *sven0925*, and extended through to the downstream Lsr2 binding site (within *sven0926*). The second construct was the same, only with the CmlR binding site and *sven0925* promoter replaced with the constitutive *ermE*\* promoter. These two reporter constructs were introduced into wild type and *lsr2* mutant strains on an integrating plasmid vector, in parallel with a promoterless plasmid control. The active *ermE\** promoter led to significantly increased β-glucuronidase activity in the wild type background relative to the CmlR-controlled promoter, suggesting reduced Lsr2 repression. In contrast, in an Δ*lsr2* background, β-glucuronidase activity did not differ significantly for the two reporter constructs, although we note that (for unknown reasons) the activity of the negative control was higher in this background (**Supplementary Fig. 3)**. Collectively, these results, when taken together with the assays described above, suggested that Lsr2 repression could be alleviated by enhancing transcription.

## DISCUSSION

Lsr2 plays a pivotal role in repressing specialized metabolism in *Streptomyces* species (4), yet it is assumed that many of these specialized metabolic clusters must be expressed under specific circumstances. Here, we probed the mechanistic basis underlying Lsr2 repression of the chloramphenicol biosynthetic cluster in *S. venezuelae*, and found that it appears to function through polymerizing along the chromosome, and bridging sites within and adjacent to the biosynthetic cluster. We further explored how Lsr2 repression was alleviated, and identified a key counter-silencing function for the cluster-situated regulator CmlR, which enhances transcription, leading to RNA polymerase effectively clearing Lsr2 from the chromosome (**Fig. 8**).

**Fig. 8:**
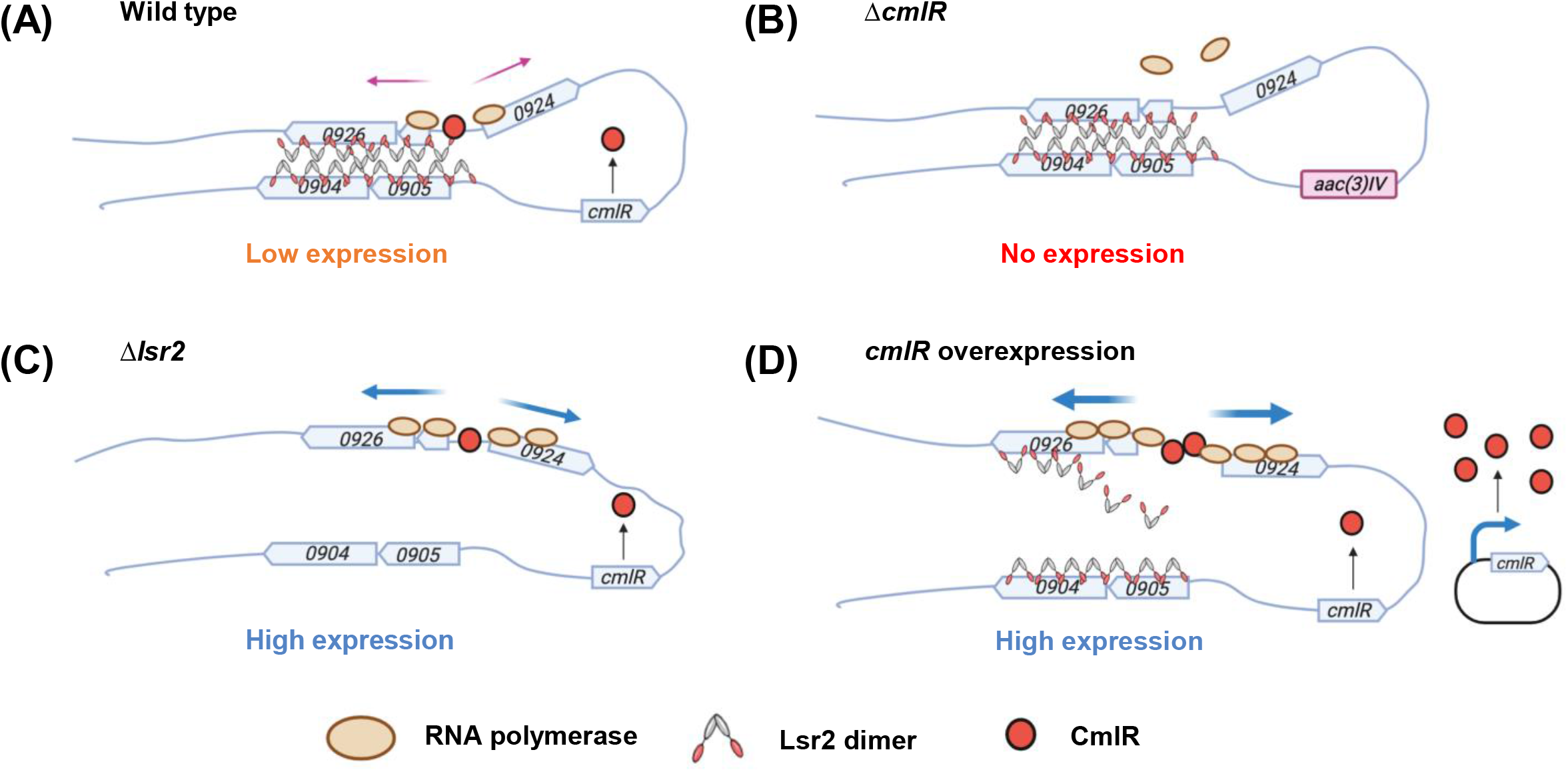
Proposed model for Lsr2 repression and CmlR counter-silencing in chloramphenicol cluster expression. **(A)** In wild type, Lsr2 represses expression of the chloramphenicol cluster by polymerizing along the chromosome and bridging sites between *sven0926* and *sven0904-0905.* Low levels of CmlR bind the divergently expressed promoter region between *sven0924* and *sven0925*, and promote baseline cluster expression and low level production of chloramphenicol. **(B)** Deleting *cmlR* leads to a complete loss of cluster expression and chloramphenicol production. **(C)** In the *lsr2* mutant, the repressing Lsr2 polymers and bridges are absent, allowing CmlR to recruit more RNA polymerase to the divergent promoter region, leading to higher cluster expression and more chloramphenicol production. **(D)** Overexpressed CmlR cooperatively binds the promoter, and its strong recruitment of RNA polymerase, and the associated increase in transcription, serves to remove Lsr2 from the chromosome and neutralizes its repressive effect, leading to high level chloramphenicol production.

Unlike most transcription factors, nucleoid-associated proteins typically bind DNA with low affinity and/or specificity, and this is consistent with our observations, where we found that CmlR bound its target sequences with far greater affinity than Lsr2. To date, the counter-silencing of nucleoid-associated protein-mediated repression has been best studied for H-NS. Three main mechanisms having been reported: (i) regulatory proteins remodel the DNA and disrupt the H-NS-DNA complex, facilitating transcription initiation by RNA polymerase (*e.g.* VirB alleviates H-NS repression at promoters of virulence genes in *Shigella flexneri*) (26,27); (ii) DNA binding proteins compete with H-NS for binding to a given site, and in doing so relieve H-NS repression (*e.g.* in *Vibrio harveyi*, the LuxR transcription factor relieves H-NS repression of bioluminescence by competing with H-NS for binding to the promoter of quorum sensing genes) (28); and (iii) transcribing RNA polymerase de-represses H-NS by remodelling or disrupting the H-NS complex, ultimately enhancing transcription [*e.g.* in *Salmonella*, PhoP reduces H-NS binding to horizontally-acquired genes by competing with H-NS for binding, and enhancing transcription by recruiting RNA polymerase (29,30)].

Counter-silencing of Lsr2 in *Mycobacterium tuberculosis* has been previously described in relation to iron metabolism (31). The expression of *bfr*, encoding a bacterioferritin, is governed both by Lsr2 and the iron-dependent transcriptional regulator IdeR. Lsr2 binds directly to the promoter of *bfrB*, thereby preventing its transcription. In iron-replete conditions, IdeR is activated by iron binding and alleviates Lsr2 repression by directly associating with the *bfrB* promoter (31). However, it is not clear whether relief of Lsr2 repression is accomplished through direct competition between IdeR and Lsr2 for binding, or by IdeR enhancing transcription levels, as appears to be the case for CmlR and Lsr2 in *S. venezuelae* (31). Counter-silencing has also been explored for the *Corynebacterium* homologue known as CgpS, using synthetic systems (32). These experiments revealed that counter-silencing of Lsr2 bound to a single site/region (*i.e.* not bridging different sequences) was most effectively achieved through competition for binding by transcription factors at the CgpS nucleation site, presumably serving to limit polymerization along the DNA (32). While it is possible that CmlR has a minor role in limiting the bounds of polymerization, our data suggest its major function is to promote transcription, and in doing so facilitate Lsr2 removal from the chromosome. What controls the expression of *cmlR* remains to be determined, as its expression is unaffected by Lsr2 activity.

Previous work has revealed that Lsr2 binding sites are found in the majority of biosynthetic gene clusters in *S. venezuelae*, including the chloramphenicol cluster (4). Our data support a model in which Lsr2 employs both an internal and external binding site to down-regulate the expression of the chloramphenicol biosynthetic genes (and production of chloramphenicol). We were curious whether such a binding configuration was associated with other Lsr2-targeted clusters. In examining the data of Gehrke et al. (4), we noted that nine clusters contained more than one Lsr2 binding site, and most of these clusters (7/9) exhibited altered transcription profiles in an *lsr2* mutant. Of the clusters associated with a single Lsr2 binding site and which also have altered transcription patterns (6 clusters), all but one has a cluster-adjacent Lsr2 binding site (within 12 genes upstream or downstream). This suggests that the model we propose for control of the chloramphenicol cluster (polymerization and bridging) may be broadly employed throughout *S. venezuelae* for repression of specialized metabolism. Whether any regulators encoded within these clusters play equivalent roles to that of CmlR in the chloramphenicol cluster, in helping to relieve Lsr2 repression, remains to be seen.

Understanding the different ways by which Lsr2 can exert its repressive effects is central to our ability to effectively manipulate its activity, and in doing so, gain access to the vast cryptic metabolic repertoire of the streptomycetes. Counter-silencing by cluster-situated activators likely represents one of many approaches employed by *Streptomyces* spp. to modulate the effects of Lsr2. It will be interesting to determine whether the activity of Lsr2 in the streptomycetes is impacted by environmental factors like H-NS (*e.g.* temperature) (19), alternative binding partners like H-NS (e.g. StpA) (33) and Lsr2 in *M. tuberculosis* (e.g. HU) (34), or post-translational modification (35,36).

## MATERIALS AND METHODS

### Bacterial strains and culture conditions

*S. venezuelae* strains were grown at 30°C on MYM (maltose, yeast extract, malt extract) agar, or in liquid MYM medium. *Escherichia coli* strains were grown at 37°C on or in LB (lysogeny broth) medium (37). *Streptomyces* and *E. coli* strains that were constructed and used are summarized in **Table 1**. Where appropriate, antibiotic selection was used for plasmid maintenance, or for screening/selecting during mutant strain construction. When assessing the importance of transcription for Lsr2 counter-silencing, *S. venezuelae* liquid cultures were grown for 16 h, after which they were exposed to the RNA polymerase-targeting antibiotic rifampicin (500 μg/mL) for 10 min.

**Table 1.** Strains used in this study.

| Strains | Genotype/characteristics/use | Reference |
| --- | --- | --- |
| ***Streptomyces*** |  |  |
| *S. venezuelae* ATCC 10712 | Wild type |  |
| E327 | *S. venezuelae* *lsr2::acc(3)IV* | (4) |
| E327A | *S. venezuelae* Δ*lsr2* | (4) |
| E332 | *S. venezuelae* *cmlR::acc(3)IV* | This work |
| E333 | *S. venezuelae* Δ*lsr2* *cmlR::acc(3)IV* | This work |
| E334 | *S. venezuelae* *sven0904_0905::acc(3)IV* | This work |
| E335 | *S. venezuelae* Δ*lsr2* *sven0904_0905::acc(3)IV* | This work |
| E336 | *S. venezuelae* Δ*lsr2* with pIJ82 carrying wildtype *lsr2* | (4) |
| E337 | *S. venezuelae* Δ*lsr2* with pIJ10706 carrying *lsr2*-3×Gly-3×FLAG | (4) |
| ***Escherichia coli*** |  |  |
| DH5α | Routine cloning | (47) |
| SE DH5α | Highly-competent (Subcloning Efficiency™) DH5α cells | Invitrogen |
| ET12567 | *dam*, *dcm*, *hsdS*, *cat* *tet*; carries *trans*-mobilizing plasmid pUZ8002 | (39) |
| Rosetta 2 | Protein overexpression host with pRARE2 which supplies 'rare' tRNAs | Novagen |

### Mutant/overexpression strain construction

In-frame deletions of *cmlR* and *sven0904-0905* were created using ReDirect technology (38). The coding sequence in cosmid 4P22 was replaced with the *aac(3)IV-oriT* apramycin resistance cassette. Mutant cosmid 4P22Δ*cmlR*::*aac(3)IV-oriT* and 4P22Δ*0904_0905*::*aac(3)IV-oriT* were confirmed using PCR, before being introduced into the non-methylating *E. coli* strain ET12567/pUZ8002 (39,40) and conjugated into wild type *S. venezuelae* or *lsr2* mutant strains. The sequence of primers used to create the disruption cassettes, and to check the integrity of the disrupted cosmids and chromosomal mutations, can be found in **Table S1**.

The *cmlR* overexpression plasmids were made by cloning the *cmlR* gene and 216 bp of its downstream sequence under the control of the constitutive, highly active *ermE\** promoter in the integrating plasmids pIJ82 and pRT801. For pIJ82, *cmlR* was amplified using primers cmlRfwd1 and cmlRrev1 (**Table S1**), digested with BamHI, and cloned into the BamHI site of pIJ82 (**Table 2**). For pRT801, *cmlR* was amplified using primers cmlRfwd2 and cmlRrev2 (**Table S1**), before being digested with SpeI and cloned into the same site in pRT801 (**Table 2**). *cmlR* presence and orientation in both plasmids were checked by PCR using vector- and insert-specific primers (**Table S1**), and construct integrity was confirmed by sequencing. The resulting plasmids, alongside empty plasmid controls, were introduced into *S. venezuelae* strains via conjugation from *E. coli* strain ET12567/pUZ8002 (**Table 1**). Strains used for chromatin immunoprecipitation (ChIP) were generated by introducing the pRT801-based *cmlR* overexpression construct into *lsr2* mutant strains complemented with either *lsr2* or *lsr2-3*×*FLAG* on the integrating plasmid vector pIJ82 and pIJ10706, respectively (**Table 1**) (4).

**Table 2.**
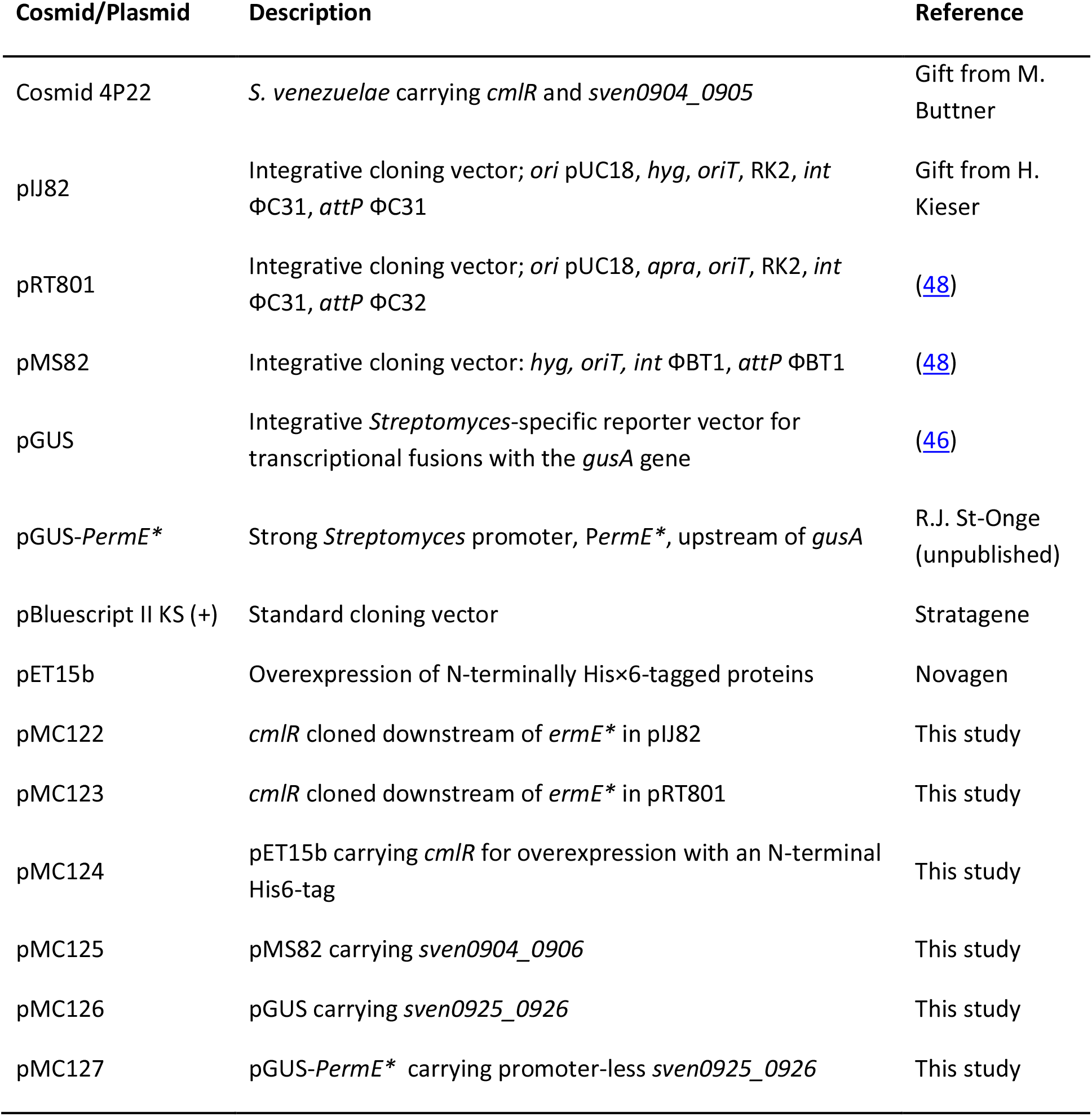
Plasmids and cosmid used in this study.

Δ*0904-0905* mutants were complemented by cloning the entire *sven0904-0906* operon, including 513 bp upstream and 123 bp downstream sequences (using primers 0904_0906CF and 0904_0906CR; **Table S1**), into the EcoRV-digested integrating plasmid vector pMS82. The resulting construct was sequenced, before being introduced into *E. coli* strain ET12567/pUZ8002, alongside empty vector control plasmids, and conjugated into *S. venezuelae* Δ*0904-0905* and Δ*lsr2*Δ*0904-0905* strains.

### Protein overexpression, purification and electrophoretic mobility shift assays (EMSAs)

Lsr2 overexpression and purification was performed as described previously (4). To overexpress CmlR in *E. coli*, the *cmlR* coding sequence was PCR amplified using primers cmlR O/E fwd and cmlR O/E rev (**Table S1**). The resulting product was digested with NdeI and BamHI, before being ligated into the equivalently digested pET15b vector (**Table 2**). After sequencing to confirm construct integrity, the resulting plasmid was introduced into *E. coli* Rosetta cells (**Table 1**). The resulting 6×His-CmlR overexpression strain was grown at 37°C until it reached an OD_600_ of 0.6, at which point 0.5 mM isopropyl-β-D-1-thiogalactopyranoside (IPTG) was added. Cells were then grown at 30°C for 3.5 h before being collected and lysed. The overexpressed protein was purified from the cell extract using nickel-nitrilotriacetic acid (Ni-NTA) affinity chromatography and was washed using increasing concentrations of imidazole (50 mM-250 mM), before being eluted using 500 mM imidazole. Finally, purified 6×His-CmlR was exchanged to storage buffer suitable for both EMSAs and freezing at −80°C (50 mM NaH_2_PO_4_, 300 mM NaCl, and 10% glycerol, pH 8).

To test Lsr2-binding specificity, EMSAs were performed using 100-222 bp probes amplified by PCR and 5’-end-labelled with [γ-32P]dATP (**Table S1**). Lsr2 (0-500 nM) was combined with 1 nM probe and binding buffer (10 mM Tris, pH 7.8, 5 mM MgCl_2_, 60 mM KCl, and 10% glycerol) in 20 μL reaction volumes. Reactions were incubated at room temperature for 10 min, followed by 30 min on ice. Any resulting complexes were then separated on a 10% native polyacrylamide gel.

To test CmlR binding to the divergent promoter region between *sven0924* and *sven0925*, a 270 bp probe encompassing the predicted binding site (amplified using primers CmlR binding F and CmlR binding R; **Table S1**) was used for EMSAs. CmlR (0-150 nM) was combined with 1 nM probe and binding buffer, as described above for Lsr2. Reactions were incubated at 30°C for 30 min, before being separated on a 10% native polyacrylamide gel. EMSA gels were exposed to a phosphor screen for 3 h, before being imaged using a phosphorimager.

### Atomic force microscopy

Lsr2 binding sites, plus considerable flanking sequences, were amplified using AFM0905F and AFM0905R (**Table S1**) for *sven0904-0905* (1612 bp product), and AFM0926F and AFM0926R (**Table S1**) for *sven0926* (2441 bp product). The resulting DNA products were cloned into pBluescript II KS(+) at the EcoRV and SmaI sites, respectively. The orientation of each fragment was assessed by PCR using vector- and insert-specific oligonucleotides (**Table S1**), and confirmed by sequencing. The resulting hybrid product was then amplified with AFM0905R and AFM0926R (**Table S1**), for use in atomic force microscopy (AFM). Lsr2 was overexpressed and purified as described above. Negative control DNA (sequences not bound by Lsr2 *in vivo*) was amplified from *S. venezuelae* genomic DNA using primers 7031F and 7031R (**Table S1**). The DNA-alone samples were prepared in 20 μL reaction volumes, and contained 0.5 nM target DNA, 10 mM Tris-HCl pH 7.6, 5 mM NiCl_2_, 40 mM HEPES, while the Lsr2+DNA samples, also prepared in 20 μL reaction volumes, contained 0.5 nM target DNA, 250 nM Lsr2, 10 mM Tris, pH 7.8, 5 mM MgCl_2_, 60 mM KCl and 10% glycerol. Different buffer conditions were used for DNA alone and Lsr2+DNA because Ni^2+^ was needed for DNA binding to the mica slide; however, it was not compatible with Lsr2 binding, so was excluded from protein-containing reactions. Reactions were incubated at room temperature for 10 min, followed by 30 min on ice. The DNA or DNA/Lsr2 was then deposited onto freshly cleaved mica surfaces (Ted Pella, Inc.) and rinsed with 1 mL nuclease-free water. Water was removed by blotting with filter paper, after which the mica surface was dried using a stream of nitrogen. AFM was performed as described in Cannavo *et al*., 2018 (41). Images (2×2 μm) were captured in air using a Bruker Bioscope Catayst Atomic Force microscope with ScanAsyst Air probes. Observed molecules were processed (through plane fit and flattening) and analysed using Image Metrics version 1.44 (42,43).

### Chromatin immunoprecipitation-qPCR (ChIP-qPCR)

ChIP-qPCR was performed as described previously (4). Strains were inoculated in 10 mL of liquid MYM medium and grown overnight, before being subcultured in duplicate, in 50 mL of MYM medium. After incubating for 18 h, formaldehyde was added to a final concentration of 1% (v/v) to cross-link protein to DNA. The cultures were then incubated for a further 30 min, before glycine was added to a final concentration of 125 mM. Immunoprecipitation of Lsr2-FLAG (or as a negative control, untagged Lsr2) was performed as described previously (44) using the FLAG M2 antibody (Sigma).

To quantify the relative abundance of target genes of interest in the ChIP DNA samples, 20 μL qPCR reactions were prepared using the LUNA^®^ Universal qPCR Master Mix (New England Biolabs), together with 2.5 μL of ChIP DNA (1:10) as template. Primer pairs used to amplify the different target DNA sequences are summarized in **Table S1**. qPCR data were analyzed using DART-PCR (45).

### Secondary metabolite extraction and liquid chromatography–mass spectrometry (LC-MS) analysis

Metabolite extraction and LC-MS analyses were performed as described previously (4), with minor modifications. Strains were grown in triplicate in 30 mL liquid MYM medium at 30°C for 2 days. Cultures were lyophilized and the resulting lyophiles were resuspended in 10 mL methanol and shaken overnight on a rotary shaker at 4°C. After centrifugation to remove particulate matter, the soluble samples were concentrated using a centrifugal vacuum evaporator (Genevac). The resulting products were then redissolved in 50% methanol and centrifuged again to remove residual particulate matter. The resulting soluble extracts were used for LC-MS analyses.

The extracts were analyzed using an Agilent 1200 LC coupled to a Bruker micrOTOF II (ESI-MS). One microliter of the injected extracts was separated on a Zorbax Eclipse XDB C18 column (100 mm × 2.1 mm × 3.5 mm) at a flow rate of 0.4 mL/min for 22 min. Extracted metabolite separation was achieved using a gradient of 0–11 min from 95% to 5% A, 11–12 min isocratic 5% A, a gradient of 12–21 min from 5% to 95% A, and 21-22 min isocratic 95% A, where A is water with 0.1% formic acid and B is acetonitrile with 0.1% formic acid. Chloramphenicol was detected using the negative ionization mode, at 321 m/z.

### β-glucuronidase (Gus) reporter assays

To test how promoter activity affected Lsr2 binding, sequences encompassing the CmlR binding site and *sven0925* promoter, through to the Lsr2 binding site in *sven0926*, were amplified and cloned into the KpnI and SpeI sites of pGUS (46) using primers 0925_26 pGUS F and 0925_26 pGUS R (**Table S1**). To replace the native promoter of *sven0925* with the constitutive *ermE\** promoter, the *ermE\** promoter was amplified from plasmid pGUS-*PermE\** (**Table 2**) using primers ermEF-X and ermER-K (**Table S1**), and cloned into the XbaI and KpnI sites of pGUS. Into the downstream SpeI site was then cloned the promoter-less *sven0925_0926* fragment amplified using 0925_26 pGUS-E*F and 0925_26 pGUS R (**Table S1**). The resulting constructs were confirmed by sequencing, and introduced into *S. venezuelae* wildtype and *lsr2* mutant strains by conjugation, alongside a promoter-less pGUS control plasmid.

The resulting pGUS-containing strains were inoculated into 10 mL MYM medium and grown at 30°C for 18 h, after which 1 mL of culture was removed and assayed for β-glucuronidase activity. Cell pellets were resuspended in lysis buffer (50 mM phosphate buffer [pH 7.0], 0.27%v/v β-mercaptoethanol, 0.1%v/v Triton X-100, 1 mg/ml lysozyme) and incubated at 37°C for 30 min. After incubation, the cell lysate was centrifuged and the resulting supernatant used in the assay. Fifty microlitres of supernatant were added to a 200 μL (total) reaction, together with the p-nitrophenyl-β-D-glucuronide substrate (PNPG; Sigma Aldrich) at a concentration of 600 μg/mL. Gus activity was determined by measuring the reaction absorbance at 420 nm, and normalizing to the OD_600_ of cultures.

## ACKNOWLEDGEMENTS

We would like to thank Lucas Koechlin for atomic force microscopy assistance, and members of the Elliot and Andres labs for helpful comments and suggestions. This work has been supported by a Canadian Institute of Health Research (CIHR) Project grant to M.A.E. (PJT-162340), and a Natural Sciences and Engineering Research Council (NSERC) Discovery grant to S.N.A. X.Z. was supported by an International Ontario Graduate Scholarship.

